# Postnatal Sensory Experience and Barrel Cortex Alterations Anticipate Autistic Traits in a Mouse Model of Cdkl5 Deficiency Disorder

**DOI:** 10.1101/2024.12.20.629347

**Authors:** Alessandra Raspanti, Riccardo Pizzo, Antonia Gurgone, Cecilia Giraudo, Giulia Sagona, Andrea Macioce, Anthony Battaglia, Lamprini Katsanou, Giorgio Medici, Francesco Ferrini, Tommaso Pizzorusso, Elisabetta Ciani, Maurizio Giustetto

**Affiliations:** “Rita Levi-Montalcini” Department of Neuroscience, University of Turin, Turin, 10125, Italy; NEUROFARBA, Department of Neuroscience, Psychology, Drug Research and Child Health, University of Florence, 50135, Florence, Italy; Department of Biomedical and Neuromotor Sciences, University of Bologna, 40126, Bologna, Italy; Department of Veterinary Sciences, University of Turin, 10095, Grugliasco, Italy; Department of Psychiatry and Neuroscience, Université Laval, Québec, QC, Canada; Scuola Normale Superiore, Biology Laboratory BIO@SNS, Pisa, 56124, Italy

## Abstract

Autistic traits may arise from atypical sensory experience during postnatal life, but whether there is a causal link between defects in cortical circuitry in the brain, altered sensory processing and social behavior remains unknown. Here, we studied tactile stimuli processing in the barrel cortex (BC) and social interactions in juvenile male mice lacking Cyclin-dependent kinase-like 5 (CDKL5), a model of a severe neurodevelopmental disease showing autistic traits and sensory impairments. We identified in these mice defects of whisker-dependent postnatal sensorimotor reflexes, NMDA receptors-dependent signaling, and dendritic orientation in thalamic inputs-receiving spiny stellate neurons. We also found that CDKL5 is required for mapping and processing whisker-derived tactile stimuli in the BC. Intriguingly, KO mice show autistic traits at p21 that are rescued by neonatal CDKL5 replacement in the BC. Our data suggest that CDKL5 is required to link tactile processing in the BC to the onset of social interaction abilities.

## INTRODUCTION

The importance of sensory experience in sculpting neuronal connectivity has been widely studied in the past years^1–3^, resulting in the definition of critical periods of brain development when experience shapes neural circuits and strongly affects the behavior^4^. Indirect evidence on the adverse effects of atypical sensory experience throughout neonatal development comes from clinical reports of individuals affected by neurodevelopmental conditions such as autism spectrum disorder (ASD)^5–8^. Among sensory modalities, somatosensory perception plays a fundamental role in early stages of brain development as it matures before other senses, beginning as early as 8 weeks into gestation, and it can strongly affect motor, cognitive and communication functions^9,10^. Interestingly, aberrant sensorimotor responses have been described in 6-month-old children followed, only in the second year of life, by cognitive and social-communication impairments^11,12^, indicating a precise temporal sequence of symptoms in ASD^13–17^. Although early somatosensory alterations are believed to be predictive of autistic traits in adulthood^11^, the mechanisms hindering the development of somatosensory circuits are still elusive, strongly limiting our understanding of ASD and the development of timely treatment options.

The whisker-mediated touch system in rodents allows tactile information to flow from the contralateral thalamus to layer 4 (L4) of the barrel cortex (BC) where thalamocortical axons (TCA) cluster into barrel-like structures, creating a precise cortical topographic map of whiskers in the snout^18–20^. This sensory neuronal network leans on precise wiring during brain development^21^ and we and others have shown that mice carrying mutation in ASD-related genes (i.e., Shank3, Fmr1, UBE3A, Mecp2 and Cdkl5) exhibit tactile abnormalities^22–29^. Patients affected by cyclin-dependent kinase-like 5 (CDKL5) deficiency disorder (CDD; OMIM: 300203^30^), a severe neurodevelopmental condition mostly caused by loss-of-function mutations of the X-linked *CDKL5* gene, display a broad spectrum of clinical signs, including early-onset epilepsy, impaired psychomotor development, abnormalities in sensory perception, and autistic-like traits^31^. We have recently shown that adult Cdkl5 KO mice, together with multiple ASD traits^32^, exhibit aberrant whisker-dependent responses that are associated with altered synaptic connectivity and activation of the barrel cortex (BC)^27,33^. Interestingly, CDKL5 expression in mice is elevated at birth, reaching the peak of expression in the cerebral cortex around p14-15^34^, when the closure of the critical period in the BC occurs^18,20^. CDKL5 is a serine/threonine kinase that participates in numerous cellular processes, including gene expression, cell division, neuronal migration, axonal outgrowth, dendritic morphogenesis, and synapse formation^35^. Moreover, the identified cytoplasmic targets of CDKL5 phosphorylation^30^, point to a major role of this kinase in the control of cytoskeletal functions underlying neuronal morphogenesis, including axonal outgrowth and polarization, dendritic branching and spines stability.

All of the above suggests that CDKL5 could contribute early during postnatal development to the sequence of experience-dependent mechanisms regulating BC maturation, tactile information representation, and social behavior. To test this hypothesis, we addressed the early causes underlying aberrant somatosensory perception in Cdkl5 KO mice and searched for the causal relationship with high-order social behavioral deficits. Thus, we investigated the developmental trajectory of sensorimotor responses in Cdkl5 KO mice during the first 15 days of life. We next assessed the integrity of thalamocortical (TC) connectivity, barrel organization, and cortical responses elicited by whisker stimulation in juvenile mutant mice (p15). Finally, we studied the role of CDKL5 on early social behavioral performances and tested the effect of its restricted re-expression in the neonatal BC on the emergence of ASD traits in Cdkl5 KO mice.

## METHODS

### Animals

All procedures were performed in accordance with European Community Council Directive 86/609/EEC for care and use of experimental animals and with protocols approved by the Italian Minister for Scientific Research (authorization D.M. n◦38/2020-PR 16/1/2020) and the Bioethics Committee of the University of Torino. Complete methods, experimental procedures and statistics are reported in the Supplementary Materials section.

## RESULTS

### Aberrant onset and development of sensorimotor responses in Cdkl5 KO pups

To assess the developmental trajectory of sensorimotor integration processes in mice lacking Cdkl5, we performed a battery of tests dependent on whisker-mediated sensorimotor responses from p3 to p15. To this aim, both Cdkl5 KO and WT littermates were evaluated in the execution of innate motor behaviors: surface righting, negative geotaxis and grasping reflexes^36^ (Supplementary Fig. 1A-C).

Surface righting. Cdkl5 KO mice did not show significant differences compared to control littermates in the latency to turn to a prone position (2-way ANOVA Genotype F (1, 245) = 1.795, p > 0.05) (Supplementary Fig. 1A).

Negative geotaxis. This test evaluates both vestibular and body coordination by assessing the ability of pups to reorient themselves towards an upwards position when placed on a downward incline. Pups lacking Cdkl5 displayed a marked inability in this test compared to WT littermates (2-way ANOVA Genotype F (1, 243) = 14.98 p < 0.001) (Supplementary Fig.1B), a sign that was particularly apparent at p15 as highlighted by Bonferroni’s multiple comparison analysis (p < 0.001).

Grasping reflexes. This test evaluates both the onset of the grasping reflex and the strength of the paw. Although the grasp reflex was detectable already at p3 in both genotypes, (Supplementary Fig. 1C), the developmental trajectory was severely impaired in Cdkl5 KO mice compared to WT pups until p12 (2-way ANOVA Genotype F (1, 243) = 18.99, p < 0.001) (Bonferroni post-hoc test: p < 0.001 p9; p < 0.05 p12).

Whisker stimulation. The repetitive presentations of unilateral whisker stimulation generate in rodent pups a startle response resulting in horizontal head movements, head-up, face or snout twitching^36,37^. Interestingly, the mild deflection of mystacial vibrissae produced clear motor responses in both WT and Cdkl5 KO pups, however, the developmental score of these physical reactions was higher in KO mice compared to WT littermates as highlighted by 2-way ANOVA (Genotype F (1, 220) = 1.982, p < 0.05) (Supplementary Fig. 1D), with no significant differences shown by the ages analyzed.

Tactile stimulation. We evaluated the writhe response to the gentle application of a tactile stimulus to different body areas including: head, back, tail and the four paws. A strong difference was seen in Cdkl5 KO pups compared to WT littermates (2-way ANOVA Genotype F (1, 245) = 13.49, p < 0.001), specifically at p15 (Bonferroni post-hoc test p < 0.001) (Supplementary Fig. 1E).

Cliff avoidance. This test relies on the mice’s inherent response to turn away from a cliff. We observed a significant impairment of Cdkl5 KO mice to avoid the cliff compared to WT littermates (2-way ANOVA Genotype F (1, 234) = 24.22, p < 0.001) (Supplementary Fig. 1F) at both p9 and p12 (Bonferroni post-hoc test p < 0.01).

Finally, we assessed if atypical sensorimotor responses shown by Cdkl5 KO pups were associated with changes in physical developmental landmarks. Body weight growth did not differ between genotypes until p15 (2-way ANOVA Genotype F (1,161) = 6. 82 p < 0.001; age F (3,161) = 282.3 p < 0.001), when a slight, but significant, decrease was observed in mutant pups. Moreover, no temporal differences in eye opening, incision eruption, and ear detachment were present between genotypes (Supplementary Fig. 2). Altogether, these results suggest that the lack of CDKL5 severely affects the physical responses underlying sensorimotor integration in early stages of life development.

### Early disruption of cortical activation, NMDA-mediated neurotransmission and intracellular signaling in the BC of juvenile Cdkl5 KO mice

Tactile sensorimotor integration in neonatal mice critically relies on the anatomical and functional integrity of the BC^38^. We found several molecular and neuronal alterations affecting this cortical area in p15 in unstimulated Cdkl5 KO mice. Our experiments show a significant reduction of c-Fos^+^ cells density, an indicator of neuronal activation, in Cdkl5 KO mice compared to WT littermates (2-way ANOVA Genotype F (1, 102) = 12.81, p < 0.001) (Fig. 1A). Interestingly, post-hoc statistical analyses revealed a selective decrease of c-Fos immunoreactivity in L4 (Bonferroni post-hoc test: L4 p < 0.01). Next, we used whole-cell patch-clamp recordings on p15 BC slices to test N-methyl-D-aspartate (NMDA)-mediated excitatory neurotransmission, as it plays a fundamental role in the development and plasticity of rodent somatosensory cortex^39^. The application of 30 µM NMDA produced an inward current (INMDA) in both WT e Cdkl5 KO BC pyramidal neurons (Fig. 1B). As we previously observed in Cdkl5 KO cortical neuronal cultures^29^, INMDA was significantly lower in Cdkl5 KO mice compared to WT littermates (WT: 822.7.4 ± 171.4.3 pA; KO: 391.2 ± 65.3 pA) (unpaired t-test p < 0.05) (Fig. 1B). We next analyzed the expression levels of NR2A and NR2B subunits in cortical extracts of WT and Cdkl5 KO juvenile mice (Fig. 1C). We found no differences in NR1 levels, the constitutive subunit of NMDAR, between WT and Cdkl5 KO (Fig. 1D) (unpaired t-test p = 0.39). In contrast, the expression of both NR2A and NR2B, the major regulatory subunits differentially expressed during BC development, were significantly lower in Cdkl5 KO cortical extracts (Fig. 1E, F) (NR2A: unpaired t-test p < 0.05; NR2B: unpaired t-test p < 0.05). Finally, we analyzed NMDAR-mediated postsynaptic signaling^40–42^ by evaluating the levels of the phosphorylated forms of Ca^2+^/Calmodulin-dependent kinase II (CaMKII), AKT and ERK in cortical synaptosomal extracts (Fig. 1G). As shown in figure 1H and I, we found a significant reduction in the levels of p-CaMKII (unpaired t-test p < 0.05) and p-AKT (S473) (unpaired t-test p < 0.05) in p15 Cdkl5 KO mice compared to WT littermates, whereas pERK did not show significant differences (Fig. 1J). No changes were detected in the total form of these proteins between genotypes (Fig. 1H-J). Altogether these data indicate that neuronal activation, NMDARs-mediated excitatory neurotransmission, and postsynaptic intracellular pathways are impaired in the BC of juvenile Cdkl5 KO mice and suggest that these alterations crucially contribute to the onset of sensorimotor deficits.

**Figure 1:**
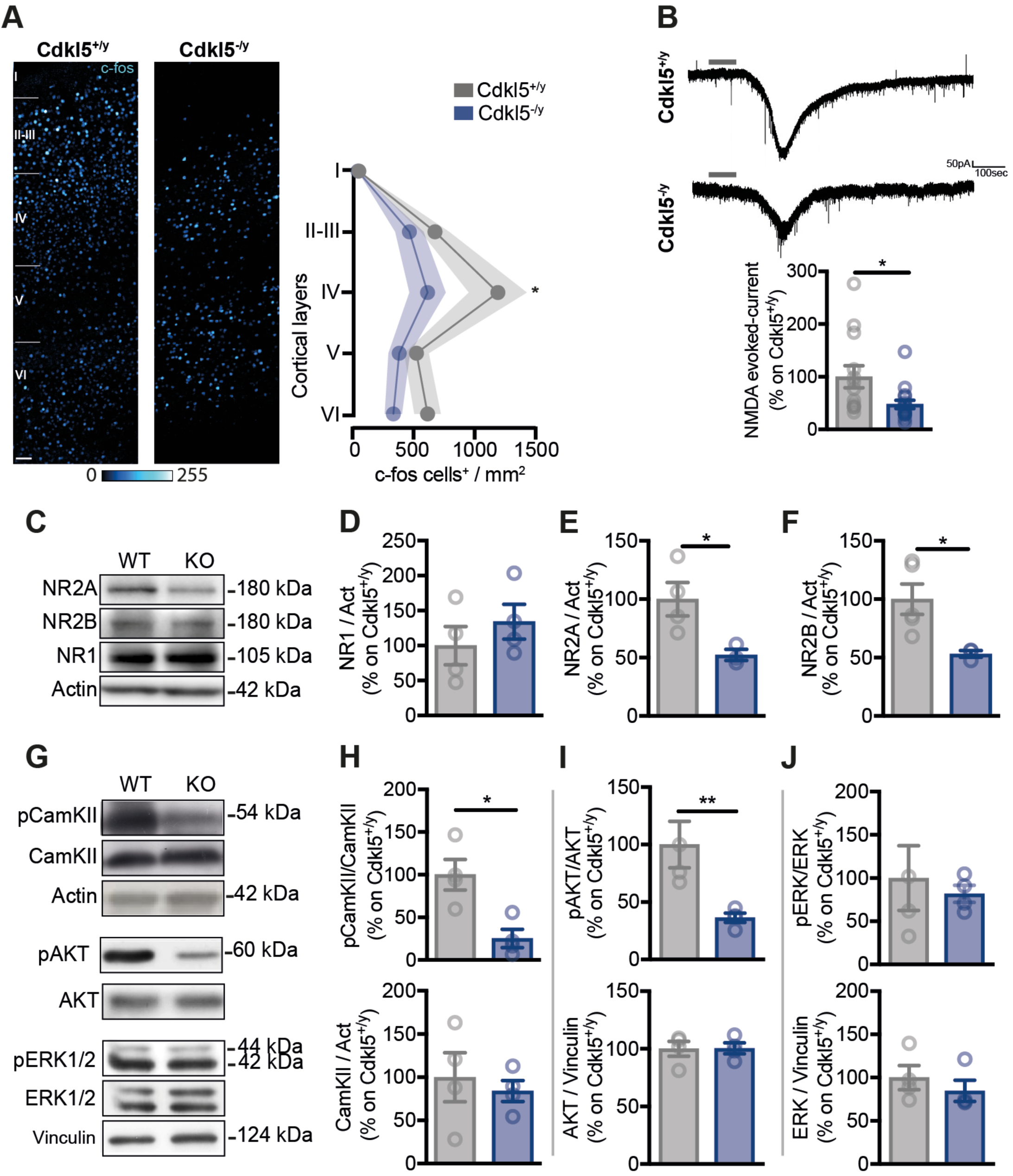
Aberrant NMDA function and NMDAR expression and dependent pathways are associated with decreased BC neuronal activation in P15 juvenile Cdkl5^-/y^ mice. (A) Representative c-Fos^+^ immunofluorescence confocal images in the BC of Cdkl5^+/y^ and Cdkl5^-/y^ mice at P15 (Scale bar 50 um) and quantification of c-Fos^+^ cells density throughout cortical layers (Cdkl5^+/y^ = 10; 9 Cdkl5^-/y^ = 9). (B) NMDA-mediated current traces and histogram showing inward current (INMDA) amplitude recorded in acute S1 slices of juvenile Cdkl5^-/y^ mice and Cdkl5^+/y^littermates following bath application of 30 µM NMDA (grey line) (cells: 13 Cdkl5^-/y^; 16 Cdkl5^+/y^). (C) Representative western blots of NR1, NR2A and NR2B in cortical extracts of Cdkl5^+/y^ and Cdkl5^-/y^ mice at postnatal at P15 and graphs showing the quantitative analysis of the total protein levels for NR1 (D), NR2A (E) and NR2B (F) levels normalized to actin (Cdkl5^+/y^ = 4; Cdkl5^-/y^= 4/3). (G) Representative western blots of pCamKII, CamKII, pERK1/2, ERK1/2, pAKT, AKT and housekeeping proteins (actin and vinculin) in synaptosomal extracts of Cdkl5^+/y^ and Cdkl5^-/y^ mice at P15. Quantitative analyses revealed a significant reduction of pCamKII/CamKII ratio in Cdkl5^-/y^ mice (H), as well as for the pAKT/AKT ratio (I) whereas no differences were identified for in pERK1/2 (J). Statistical analyses: t-test, 2-way ANOVA followed by Bonferroni post-hoc test *p < 0.05, **p < 0.01.

### The loss of Cdkl5 interferes with dendritic asymmetry of L4 spiny stellate cells

The establishment of TC wiring requires NMDARs-mediated neurotransmission19, which we have found altered in the BC of Cdkl5 KO mice. Because spiny stellate neurons (SSNs) in L4 are the main recipient of TC inputs, relaying tactile information from whiskers in the snout to the BC^43^, we assessed the integrity of their morphology and connectivity. To this aim, AAV-mediated retrograde labeling of TCA was obtained by injections in the BC of both WT and KO mice, and images were captured in the caudal striatum of p15 mice by confocal microscopy (Fig. 2A-B). Interestingly, TCA organization did not reveal substantial differences in axonal density between WT and Cdkl5 KO mice (Fig. 2C) (unpaired t-test p < 0.05). To exclude any difference due to AAV infection, we normalized TCA density values on the mean EGFP intensity shown by the injection site in the BC. As shown in Fig. 2D, also after this normalization no differences were found between genotypes, thus suggesting the integrity of TCA extension is preserved in juvenile Cdkl5 KO mice. Finally, we assessed the intensity of EGFP-expression in TCA across the area of interest (dashed area in Fig. 2B). This analysis revealed a general reduction of TCA fluorescence in KO mice, suggesting that lack of Cdkl5 may alter either retrograde transport mechanisms or axonal size (Fig. 2E) (2-way ANOVA Genotype F (1, 6264) = 398.7, p < 0.001). To exclude major anatomical abnormalities of the thalamic VPM nucleus, we assessed its cellular density (Supplementary Fig. 3). The analysis of DAPI^+^ cells density revealed no difference between genotypes.

**Figure 2:**
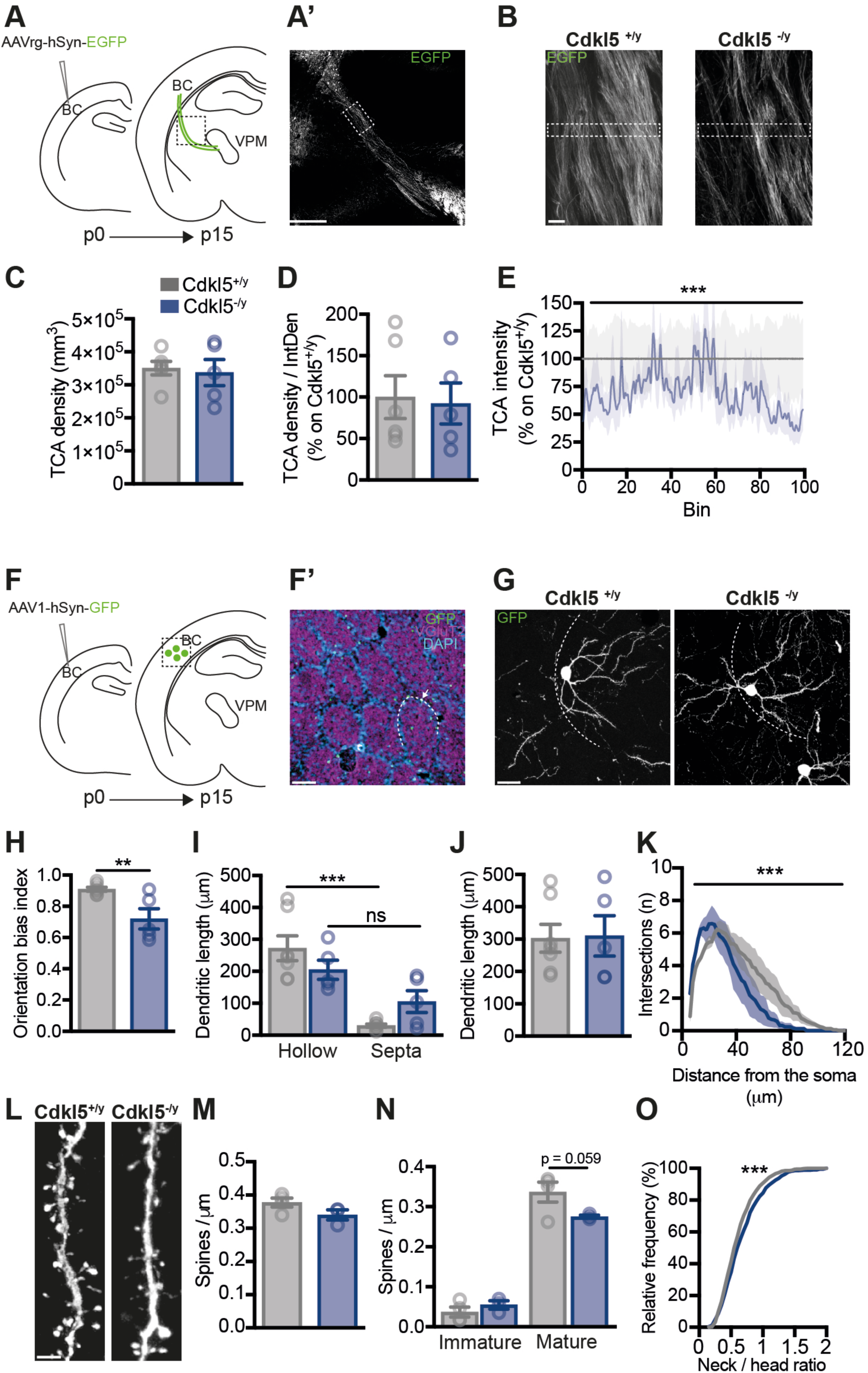
Cdkl5 loss impacts dendritic asymmetry of L4 spiny stellate cells. (A) Schematic illustrating stereotaxic AAVrg-hSyn-EGFP injections into the BC of p0 Cdkl5^+/y^ and Cdkl5^-/y^ mice to label TC projections to BC and (A’) confocal micrograph showing labeled TCAs of p14 mice in the boxed area in panel A. B) Higher magnification confocal images of the outlined area in panel A’ in p14 Cdkl5^+/y^ and Cdkl5^-/y^ mice. (C, D) The analysis of the density of individual EGFP^+^ TCAs (C) and the TCAs density corrected on the mean intensity of the injection site in the BC (D) did not reveal any significant differences between Cdkl5^+/y^ and Cdkl5^-/y^ mice. (E) The analysis of the TCA intensity quantified in the area enclosed by the dashed line in B shows a reduction in the Cdkl5^-/y^ mice suggesting a decreased TCA diameter. F) Schematic representation of the sparse stereotaxic AAV-hSyn-GFP injections into the BC of p0 Cdkl5^+/y^ and Cdkl5^-/y^ mice to label spiny stellate cells (SSNs) and (F’) confocal micrograph on tangential sections of flattened cortices showing labeled SSNs in L4 BC (boxed area in panel F) identified by VGluT2 staining in p15 mice. Arrow indicates a L4 SSN whose soma is located in the barrel wall (dotted line). (G) Higher magnification confocal images of flattened cortices showing GFP^+^ L4 SSN of p15 Cdkl5^+/y^ and Cdkl5^-/y^ mice located in the barrel wall (dotted line). (H-J) The orientation bias index (H) (i.e. the ratio of the hollow projecting dendritic length to total dendritic length) was significantly decreased in Cdkl5^-/y^ SSNs; Cdkl5^-/y^ SSNs showed a trend towards a decrease in hollow-projecting dendritic length and a trend towards an increase in septa-projecting dendritic length (I). (J) Total dendritic length is unchanged in Cdkl5^-/y^ SSNs. (K) The Sholl analysis of SSN dendritic branching revealed a reduced number of intersections in Cdkl5^-/y^ in comparison to Cdkl5^+/y^ (Cdkl5^+/y^ = 5; Cdkl5^-/y^ = 4; 6 SSNs/animal). (L) Micrographs depicting secondary dendrites from Cdkl5^+/y^ and Cdkl5^-/y^ SSNs. (M) No statistically significant differences in dendritic spine density were observed in Cdkl5^-/y^ compared to Cdkl5^+/y^ mice. (N) A trend towards reduction was found in the density of mature dendritic spines in Cdkl5^-/y^ compared to Cdkl5^+/y^ SSNs (N); cumulative analysis of spine density based on neck/head ratio showed a statistically significant decrease between the 2 experimental groups (O) (Cdkl5^+/y^ = 4; Cdkl5^-/y^ = 3; 5 SSNs / animal; 2 secondary dendrites / SSN). Scale bars: A’, F’ 100 µm; B, G 20 µm; L, 2 µm. Statistical analysis: t-test, 2 way-ANOVA, Bonferroni’s post hoc test, Mann-Whitney, ** p < 0.01, *** p < 0.001.

Next, we studied the morphology of SSNs, the postsynaptic targets of TCA inputs. Barrels form in the first week after birth once they are reached by presynaptic TCA contacting L4 SSNs that are localized at barrel edges and become polarized by extending dendrites asymmetrically toward barrel hollow upon receiving TCA inputs^18,44,45^. Thus, we assessed the role of CDKL5 on SSNs polarity by investigating the dendritic organization of sparsely-labeled GFP^+^ SSNs in tangential sections of flattened cortices of mice injected with AAV-hSyn-GFP in the BC at p0 (Fig. 2F)^46^. GFP^+^ neurons, whose cell bodies were located at barrel edges (Fig. 2F’, G), were selected and the orientation bias index (OBI) of their dendrites, i.e., the ratio of the length of hollow-projecting dendrites to the total dendrite length, was determined. Intriguingly, whereas in WT mice SSNs extended most dendrites toward the hollow, Cdkl5 KO SSNs showed numerous dendritic segments oriented into the septa (Fig. 2G). This qualitative observation was confirmed by quantitative evaluation of both OBI (unpaired t-test p < 0.05) and total dendritic length, measured in barrel hollows and septa (2-way ANOVA followed by Bonferroni, hollow vs septa, WT p < 0.001, KO p = 0.08) (Fig. 2H, I). The atypical orientation shown by Cdkl5 KO SSNs did not result from abnormal dendritic growth as the total dendritic length was similar between genotypes (Fig. 2J) (unpaired t-test p = 0.92). To characterize whether the loss of CDKL5 might also impact the branching complexity of SNNs dendrites, we conducted a Sholl analysis. As shown in fig. 2K, the 2-way ANOVA revealed a statistically significant difference between genotypes (p < 0.001) with the KO SSNs showing a reduced number of intersections compared to WT cells.

We previously showed that CDKL5 is necessary for dendritic spine maturation and stabilization in cortical excitatory neurons located in S1^47^. Given the hierarchical organization of sensory inputs processing in the BC, with SSNs being the primary postsynaptic target of TCA^48^, we assessed dendritic spines structure decorating these neurons. We found that the density of dendritic spines protruding from secondary dendrites of GFP^+^ SSNs did not differ between genotypes (Fig. 2L,M). In contrast, when we evaluated their morphology, 2-way ANOVA revealed a genotype x spine category (i.e. mature vs. immature) interaction (F (1, 10) = 5.31, p < 0.05), although Bonferroni post-hoc test did not detect significant differences between genotypes (immature: p = 0.73, mature: p = 0.059) (Fig. 2N). On the other hand, cumulative distribution of spine density based on neck/head ratio values confirmed these structural alterations by revealing a statistically significant difference (Fig. 2O) (Mann-Whitney, p < 0.001). Thus, these data indicate that Cdkl5 loss produces an atypical orientation of SSNs dendrites and alters the maturation of the dendritic spines decorating these cells. These findings show that Cdkl5 loss, while it seems to only mildly affect TCA growth and path-finding into the BC, produces severe morphological alterations in L4 SSNs that can underlie abnormal thalamocortical connectivity.

### Cdkl5 KO mice show early atypical TC connectivity in L4 of the BC

Tangential sections through L4 of the BC were immunostained with an antiserum against the vesicular glutamate transporter 2 (VGluT2), to selectively label TCA terminals^27^. Although the outline of TCA in the posteromedial barrel subfield, where mystacial vibrissae are represented, did not show any substantial difference between genotypes (Fig. 3A), a significant decrease of VGluT2 immunofluorescence was observed in Cdkl5 KO mice compared to WT littermates (unpaired t-test p < 0.05) (Fig. 3B), indicative of altered synaptic integrity. In agreement with this, we found a robust reduction in the density (Fig. 3D), size (Fig. 3E), and fluorescence intensity (Fig. 3G) of VGluT2^+^ puncta in Cdkl5 KO mice compared with WT littermates (unpaired t-test p < 0.001). In addition, Cdkl5 null mice showed lower density of VGluT2-Homer1bc^+^ puncta appositions (unpaired t-test p < 0.001), resulting in a greater number of postsynaptic spines lacking an identifiable thalamic input (Fig. 3F). Interestingly, both density and size of Homer1bc^+^ puncta did not differ between genotypes (unpaired t-test p > 0.05) (Fig. 3H, I). SSNs in L4 make recurrent excitatory intracortical connections with neighboring SSNs that are needed to amplify cortical activation during sensory stimulation^21^. We found that in Cdkl5 KO mice the integrity of such intracortical connectivity was preserved. Both density and size of VGluT1^+^ puncta were unaffected in Cdkl5 KO mice compared to WT littermates (unpaired t-test p > 0.05) (Fig. 3K-M), as was the density of VGluT1-Homer^+^ juxtapositions (unpaired t-test p > 0.05) (Fig. 3L). These results indicate that at this developmental stage, by affecting the growth of TCA presynaptic terminals, the lack of CDKL5 tampers with the formation/stabilization of axospinous TC synapses in L4 of the BC.

**Figure 3:**
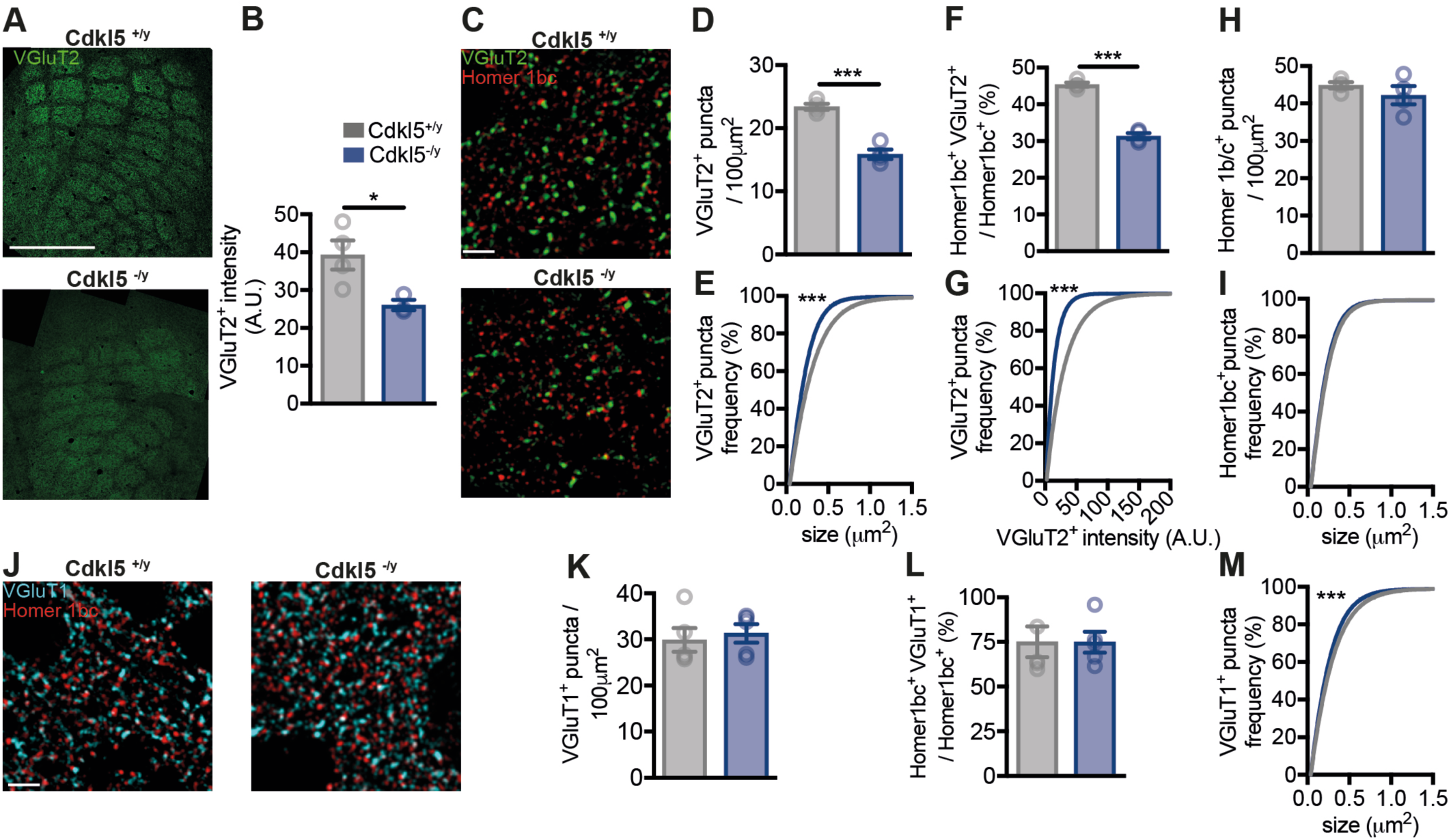
Cdkl5 loss induces circuit-specific defects of L4 barrel cortex connectivity in juvenile Cdkl5^-/y^ mice. (A) Representative VGluT2 immunofluorescence of tangential sections through L4 revealing the organization of thalamo-cortical (TC) afferents in the BC of both Cdkl5^+/y^ and Cdkl5^-/y^ mice (scale bar 500 µm) and (B) histogram showing the analysis of VGluT2^+^ staining intensity (n = 4). (C) Micrographs showing TC synapses in L4 of the BC from Cdkl5^+/y^ and Cdkl5^-/y^ mutants (scale bar 5 µm). (D-G) Quantification of VGluT2^+^ TC inputs density (D), size (E), intensity (G) and the percentage of VGluT2^+^ TC afferents juxtaposed to a Homer1bc^+^ puncta (F) (n = 4) revealed a strong impairment of TC synapses in the Cdkl5^-/y^ mice compared to Cdkl5^+/y^ littermates. (H,I) No overall changes in either the density or the size of Homer1bc^+^ dendritic spine were identified. (J) Micrographs showing the cortico-cortical CC (F) synapses in L4 of the BC from Cdkl5^+/y^ and Cdkl5^-/y^ mutants (scale bar 5 µm). (K-M) Quantification of the density of VGluT1^+^ puncta (K) as well as the percentage of VGluT1^+^ afferents juxtaposed to a Homer1bc^+^ puncta (L) revealed no overall changes in the number of intra-cortical excitatory terminals between Cdkl5^+/y^ and Cdkl5^-/y^. Nevertheless, mutants exhibit a decrease in the VGluT1^+^ puncta size (M) (Cdkl5^+/y^ = 4; Cdkl5^-/y^ = 5). Statistical analysis: t-test, Mann-Whitney, *p < 0.05, *** p < 0.001.

### Enhanced cortical response to whisker stimulation in juvenile Cdkl5 KO mice

To functionally assess sensory information processing within the BC, we examined whisker-evoked responses using intact skull intrinsic optical signal (IOS) imaging of the BC in juvenile animals (Fig. 4A)^49^. Figure 4B shows typical examples of responses consisting in a decrease in reflectance (darker area) induced by tactile stimulation in the BC. Data quantitation showed that mechanical stimulation of the contralateral mystical vibrissae produced similar amplitude of the responses in the two genotypes (unpaired t-test p = 0.12) (Fig. 4C). To elucidate whether the BC hyperactivation might be associated with an altered segregation of whisker-mediated inputs into corresponding L4 receptive fields, we employed the novelty exposure test, as described in^50^. In this task, we assessed c-Fos expression in the C2 barrel corresponding to the stimulated vibrissa as in (Fig. 4D). In agreement with the IOS results, we reported an increase in c-Fos intensity in the BC area corresponding to the remaining whisker C2 (Fig. 4E,G) (unpaired t-test, p < 0.05) as well as a trend towards an increase in the density of c-Fos^+^ cells (Fig. 4F) (unpaired t-test, p=0.11). To evaluate whether the whisker-related information is confined to its corresponding cortical barrel in the Cdkl5 KO, we analyzed the c-Fos immunofluorescence in C2 and neighboring barrels^51^ (Box areas in Fig. 4E). The intensity distribution analysis revealed that in Cdkl5 KO mice c-Fos immunosignal was not restricted to the intact C2-whisker barrel rather appeared more widespread in neighboring barrel regions compared to WT animals (2-way ANOVA Genotype: F (1, 2000) = 35.80, p < 0.001) (Fig. 4H). These findings indicate that Cdkl5 KO mice exhibit a sensory-driven hyperactivation of the BC that is paralleled by altered mapping of whisker activity.

**Figure 4:**
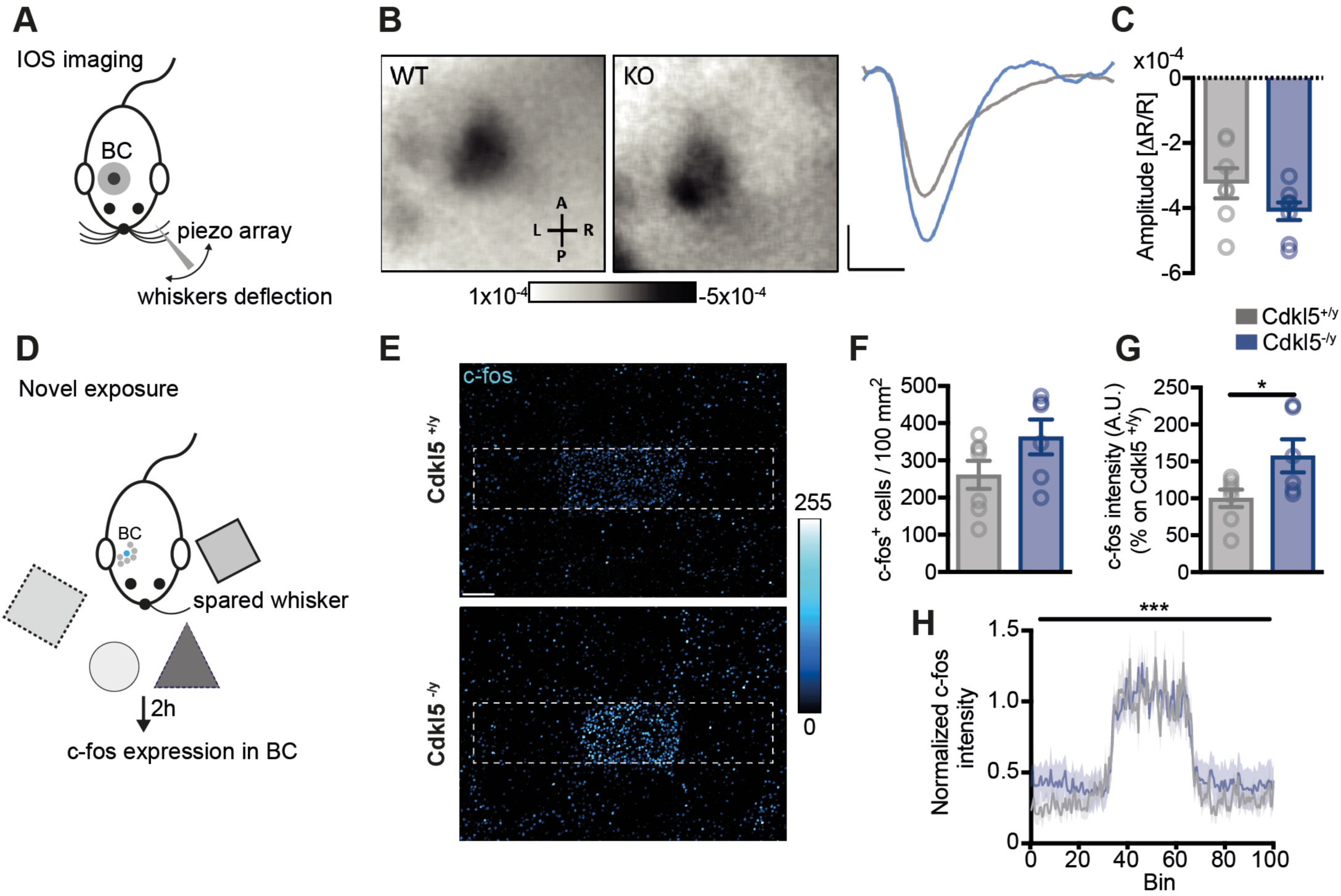
Cdkl5^-/y^ mice show aberrant cortical response following whisker stimulation. (A) Schematic of the experimental setup of the intrinsic optical imaging (IOS). 4Hz whiskers deflection in the left whisker pad was produced by using a piezo-motor based stimulator. (B) Responsive area of contralateral BC of Cdkl5^+/y^ and Cdkl5^-/y^ mice after whisker stimulation and related traces. (C) Quantification of IOS amplitude revealed an increased responsiveness of BC in Cdkl5^-/y^ mice despite not reaching the significance (p = 0.12) (Cdkl5^+/y^ = 7; Cdkl5^-/y^ = 8). (D) Schematic for the novelty exposure paradigm for assessing whisker-dependent neuronal activity. Mice were cut all but the C2 whisker and then placed in a novel texture environment. Whisker-evoked activity in the barrel cortex was assessed 2 hours after novelty exposure by c-Fos^+^ expression. (E) Representative images of tangential sections for c-Fos^+^ immunofluorescence intensity in the C2 barrel of Cdkl5^+/y^ and Cdkl5^-/y^ mice subjected to the novelty exposure paradigm. (F, G) Quantification histograms of c-Fos^+^ cells density and intensity in the corresponding C2 barrel showed, in the mutant mice, cortical hyperactivation as indicated by the trend toward increase of c-Fos^+^ cells (F) and the significant increase in c-Fos^+^ intensity (G). (H) Graph showing the profiles of c-Fos^+^ expression intensity across the ROIs outlined in (E); c-Fos^+^ intensity is normalized to the peak C2 barrel intensity (Cdkl5^+/y^ = 7; Cdkl5^-/y^ = 8). Scale bar: 100 µm. Statistical analyses: t-test, 2-way ANOVA *p < 0.05, ***p < 0.001.

### Autism-like behaviors appear early in Cdkl5 KO mice

Early tactile stimulation and experience during development are essential for brain development, cognition, and adult social behaviors^9,52–54^. To assess whether the alterations on sensorimotor responses, TC connectivity, and BC activation produced by the lack of Cdkl5 are associated with the emergence of ASDs signs, we performed a battery of behavioral tests (i.e., three-chamber sociability test, juvenile play, and repetitive behavior) from p21 to p25, a time-window when the first social interactions between conspecifics occur^55^.

First, we employed a three-chamber social arena to test the Cdkl5 KO mice for self-initiated interactions with a WT stranger mouse. At this age, Cdkl5 KO mice show no interest in the social chamber, as revealed by the increased time spent in the side of the arena with the empty cup (Bonferroni post-hoc test: WT p > 0.05; KO p < 0.05). For both WT and KO mice, the animals spent significantly less time in the central chamber than in the others, revealing the absence of anxious behavior (Bonferroni post-hoc test p < 0.01) (Fig. 5A,B). Moreover, we calculated the Sociability Index (SI) to allow the direct comparison of social behavior of each group as in^56^ (Fig. 5C). This analysis highlighted that, despite the low social preference displayed by juvenile WT mice, Cdkl5 KO mice show an SI below the 0, indicating a reduced social interest (unpaired t-test p < 0.05). Mice were next tested for direct social interaction with the juvenile play test (Fig. 5D). In agreement with three-chamber data, Cdkl5 KO pairs displayed reduced interactions compared to WT pairs (unpaired t-test p < 0.05) (Fig. 5E). To exclude that impaired locomotion contributes to atypical social behaviour displayed by Cdkl5 KO mice, we performed the open field test. As shown in figure 5H-K, juvenile KO mice show increased explorative behavior, increased mean speed, and a strong decrease in resting time (unpaired t-test p < 0.01), indicating that enhanced explorative behavior, that we and others showed in adult Cdkl5 KO mice^29^, is already detectable in early developmental stages. Finally, we assessed repetitive behaviors, another core sign of ASDs, in juvenile mice in a home-cage environment. Cdkl5 KO mice showed significantly increased time engaging in stereotypic behaviors such as jumping and digging compared to littermate controls (unpaired t-test p < 0.05) (Fig. 5F-G). Therefore, these data indicate that Cdkl5 KO mice exhibit early-onset atypical spontaneous social interactions compared to WT littermates confirming that defective morpho-functional organization of the BC is paralleled by the onset of autistic-like behaviors.

**Figure 5:**
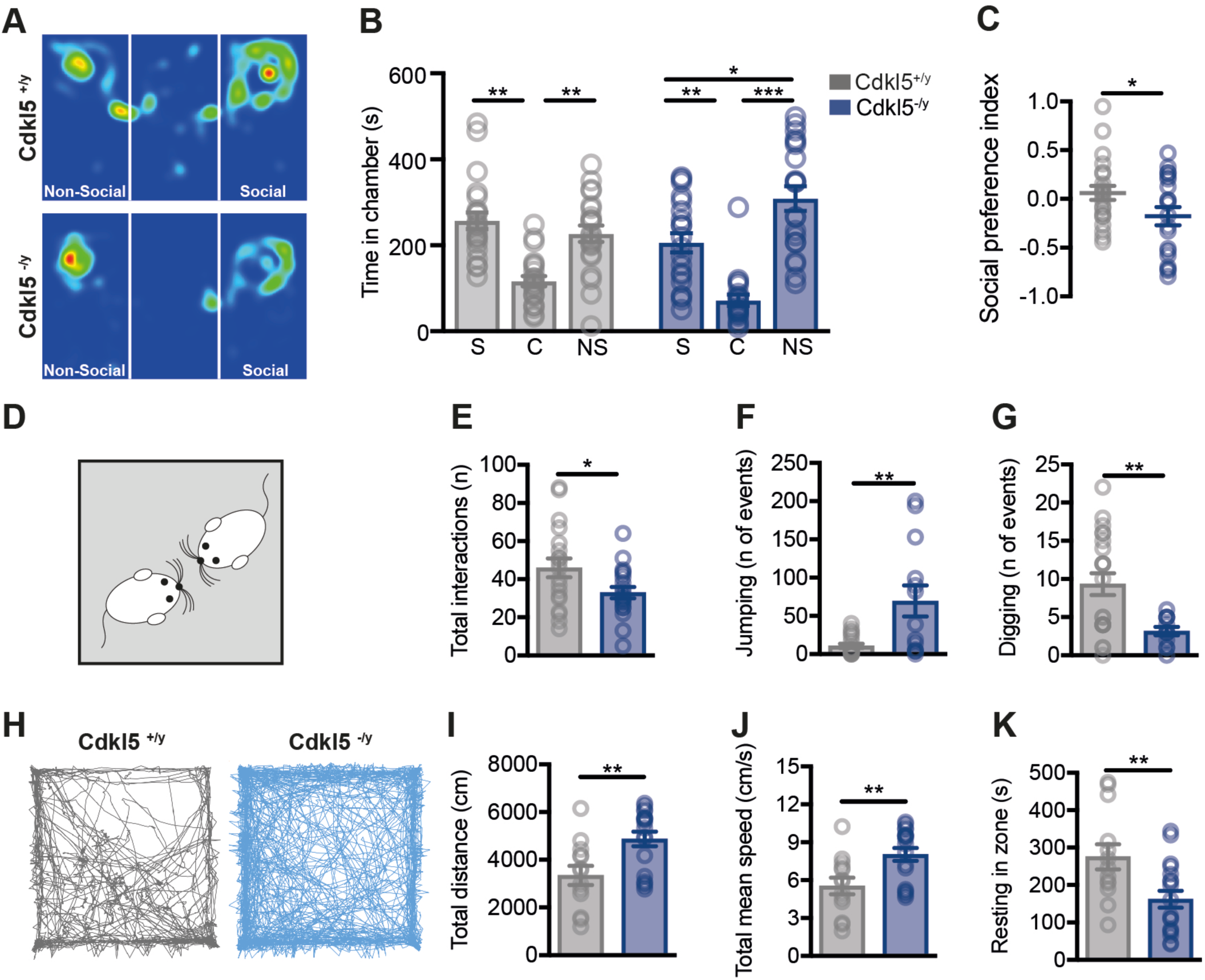
Juvenile P21-P25 Cdkl5^-/y^ mice show autistic-like phenotypes. (A) Representative heat map images of Cdkl5^+/y^ and Cdkl5^-/y^ mice in the three-chamber test. (B,C) Quantification of the time spent in the social (S), center (C) and non-social (NS) chamber (B) and the preference index in social exploration (C) (calculated as follow: if social is S and non-social is NS, then index=(S-NS)/(S+NS) revealed decreased social interaction in the mutant mice (Cdkl5^+/y^ = 22; Cdkl5^-/y^ = 20). (D,E) Schematic of the juvenile play test (D) and quantification of the reciprocal social interaction events (nose-to-nose interaction, body sniffing, anogenital sniffing, follow, push-crawl) (E) which showed a reduction in Cdkl5^-/y^ pairs compared to Cdkl5^+/y^ pairs (Cdkl5^+/y^ = 14; Cdkl5^-/y^ = 16). (F,G) Increased stereotypical behaviors in juvenile Cdkl5^-/y^ mice as revealed by the number of jumping (F) and digging (G) events (Cdkl5^+/y^ = 12; Cdkl5^-/y^ = 10). (H-K) Increased explorative behavior in juvenile Cdkl5^-/y^ mice is evaluated by the open field test. (H) Representative trajectories and quantification of (I) total distance moved, (J) speed and total resting time pointed out an early hyperactivity of Cdkl5^-/y^ mice compared to Cdkl5^+/y^ littermates (Cdkl5^+/y^ = 13; Cdkl5^-/y^ = 18). Statistical analyses: t-test, 2-way ANOVA followed by Bonferroni’s post hoc test, *p < 0.05 **p < 0.01 ***p < 0.001.

### Neonatal correction of Cdkl5 expression in the BC alleviates ASD-like behaviors

To address whether the onset of ASD-like traits in Cdkl5 KO mice are directly produced by the abnormalities affecting BC connectivity and responsiveness, we evaluated the ability of Cdkl5 re-expression, by employing a gene replacement strategy^57^, in the BC to correct the ASD-like deficits. To this aim, AAVPHP.B_CDKL5 vector or vehicle was bilaterally administered in the BC of Cdkl5 KO and WT mice, respectively, at p0 (Fig. 6A). Mice were tested for social and repetitive behaviors 24 days after the injection. In agreement with what we observed in untreated mice, vehicle-injected Cdkl5 KO pairs exhibited an almost significant decrease in social interaction events compared to WT mice (1-way ANOVA, Sidak post test; WT-vehicle vs KO-vehicle, p = 0.06) (Fig. 6B). Intriguingly, the correction of CDKL5 expression restricted to the BC produced a complete correction of atypical social behavior as indicated by the number of spontaneous social interactions displayed by AAVPHP.B_CDKL5-administered Cdkl5 KO mice (1-way ANOVA, Sidak post test, KO-vehicle vs KO-AAV, p < 0.05) (Fig. 6B). On the other hand, CDKL5 re-expression led to partial improvement of repetitive behaviors (Fig. 6C,D). Vehicle-administered KO mice displayed atypical jumping (1- way ANOVA, Sidak post test; WT-vehicle vs KO-vehicle, p = 0.07) and digging phenotypes (1-way ANOVA, Sidak post test; WT-vehicle vs KO-vehicle, p < 0.05), in line with our previous observations (Fig. 6F,G). In contrast, while the injection of AAVPHP.B_CDKL5 had no effects on the number of jumping events (1-way ANOVA, Sidak post test; KO-vehicle vs KO-AAV, p = 0.50; WT-vehicle vs KO-AAV, p < 0.01) (Fig. 6C), the correction of CDKL5 expression in the BC completely rescued digging behavior in Cdkl5 KO mice (1-way ANOVA, Sidak post test, KO-vehicle vs KO-AAV, p < 0.05; WT-vehicle vs KO-AAV, p = 0.51) (Fig. 6D). To confirm the efficiency of gene transfer by the AAVPHP.B_CDKL5, CDKL5 expression was assessed by fluorescence in-situ hybridization using a probe designed on the CDKL5 exon 4 as in^57^. As shown in Fig. 6E, the re-expression of CDKL5 was limited to the BC. Therefore, these data indicate that early correction of Cdkl5 expression in the BC of Cdkl5 KO mice is able to alleviate early onset spontaneous social interaction defects as well as to correct, at least partially, stereotypical behaviors, thus disclosing a causal link between CDKL5 function in the BC and early onset autistic-like behavior.

**Figure 6:**
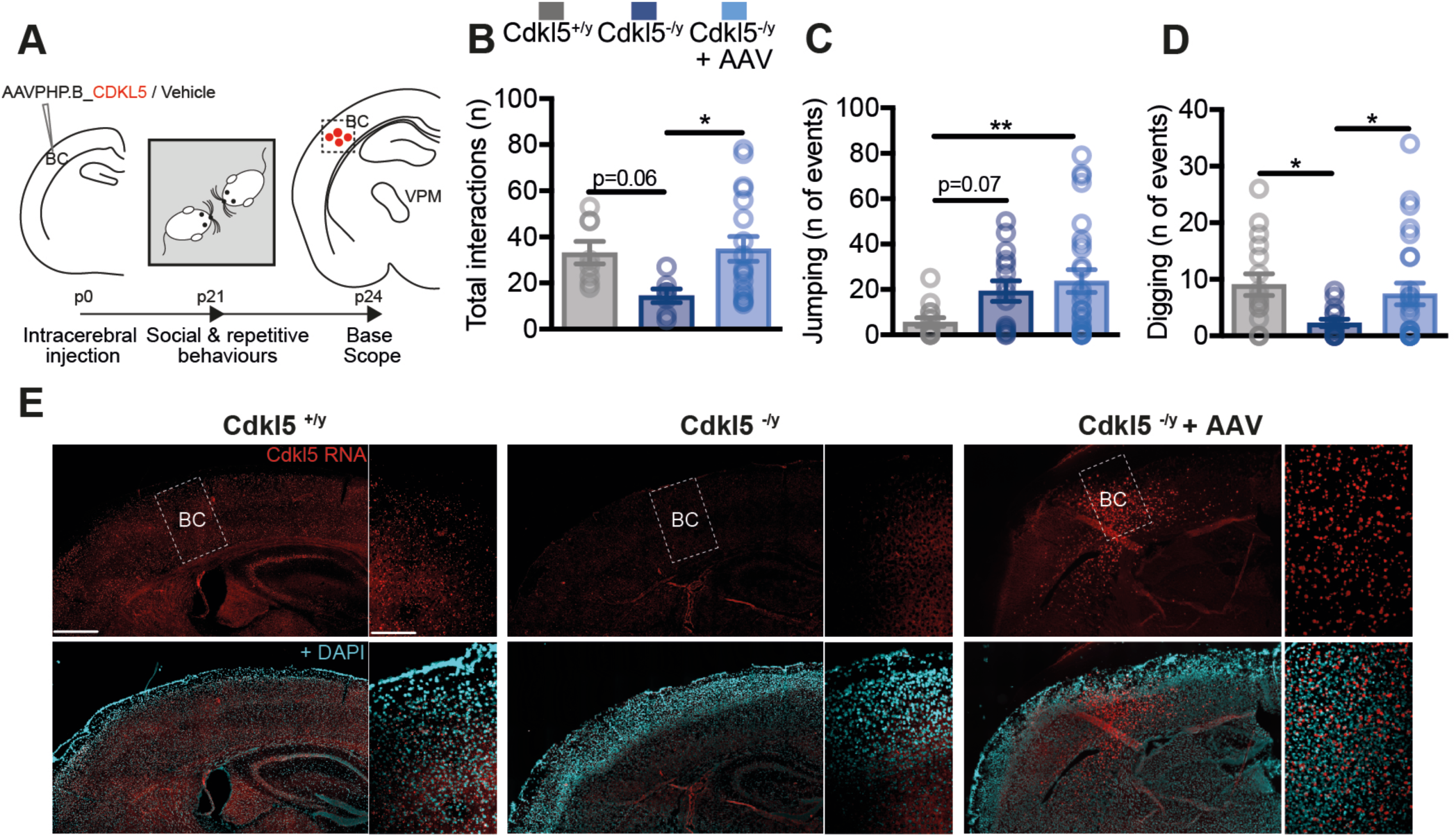
Neonatal CDKL5 expression in KO mutant mice reverts ASD-like symptoms in juvenile mice. (A) Schematic representation of neonatal CDKL5 re-expression. Mice underwent bilateral injection with AAVPHP.B_CDKL5, at a dose of 1*10^12^ (100nL), in BC at P0. Mice were tested for both social and repetitive behaviours 3 weeks later; finally, brains were collected for analysing CDKL5 re-expression by RNAScope. (B) Restored levels of social behavior in juvenile Cdkl5^-/y^ mice 3 weeks after AAV9 injection as indicated by the analysis of the number of direct interactions (vehicle injected Cdkl5^+/y^ = 24; vehicle injected Cdkl5^-/y^ = 18; AAV injected Cdkl5^-/y^ = 19). (C,D) Impact of CDKL5 re-expression on stereotypical behaviours exhibited by Cdkl5^-/y^ mice. No significant effects were observed in the jumping events (C) whereas digging impairments were completely corrected by AAV injection (D). (E) Representative confocal images of CDKL5 mRNA detected by RNA Scope in the BC of Cdkl5^+/y^ and Cdkl5^-/y^ mice injected with vehicle, and of Cdkl5^-/y^ mice injected with the AAVPHP.B_CDKL5 vector. The dotted areas outline the BC and are shown as higher magnification panels. Sections are counterstained with DAPI. Scale bars: 30 µm. Statistical analysis: 1-way ANOVA followed by Sidak post-hoc test: *p < 0.05 **p < 0.01.

## DISCUSSION

The ability to plan and execute an appropriate motor response to environmental demands is based on the proper development of the nervous system, through which innate sensorimotor reflexes, elicitable in humans within the first six months of life^58^, are progressively inhibited and replaced by postural reflexes^59,60^. If primary reflexes persist beyond the typical developmental time frame, they can interfere with maturation processes and diminish the brain’s capacity to process sensory information efficiently^61–63^.

With this study, we provide evidence that mice lacking CDKL5 show a delay in the disappearance of innate sensorimotor reflexes, especially those based on whisker-BC mediated processing (i.e., negative geotaxis test and tactile stimulation reflex), remaining present at p15. This observation aligns with recent findings showing that certain reflexes persist beyond the typical developmental timeline in children with ASD, while others do not emerge when expected^64^, and support the idea of monitoring sensorimotor reflexes as early autism biomarkers. Such behavioral development is accompanied by the formation of the barrel system in the S1 cortex, that is responsible for whisker-dependent sensorimotor control and devised as an integrated and refined network connecting whiskers, thalamus, and cortex. Indeed, we show for the first time that CDKL5 is required for the early developmental processes shaping the organization of the BC. First, we show that the reduced efficacy of NMDA-mediated transmission in the BC of juvenile Cdkl5 KO mice produces alterations in signaling pathways (i.e., CaMKII and AKT) crucial for the formation, maintenance and plasticity of synaptic connectivity^65^. Moreover, the number of c-Fos^+^ cells in L4 are robustly reduced in the BC of Cdkl5 KO mice at p15, indicating that abnormal cortical activation shown by adult KO mice^27,29,66^ originate early. Intriguingly, our data disclose that CDKL5 positively regulates NMDARs expression and function during the critical developmental periods for whisker map formation. By showing that both _I_NMDA and expression of NR2A and NR2B subunits are reduced in the absence of CDKL5 starting from p15, these experiments are in agreement with our previous data obtained in primary cultures of S1 neurons from Cdkl5 KO mice^29^. Although previous studies investigated the involvement of CDKL5 in NMDARs function, no conclusive evidence is available. In a full KO mouse model, CDKL5 loss was accompanied by defective NR2B amount in hippocampal PSD fraction^67^, whereas no changes were detected in both hippocampal and cortical membrane-bound levels of both NR2A and NR2B in a Cdkl5 KI (R59X) model^68^. The most parsimonious explanation of these differences arises from either the CDKL5 mouse model used or the cellular fractions investigated in these studies. Consistently with our current findings, previous studies show that cortex-specific deletion of NMDAR subunits tampers with the integrity of BC organization and SSNs dendritic orientation^69–71^, cellular signs that we report in Cdkl5 KO animals. In addition, metabotropic glutamate receptor 5 (mGluR5) activity was shown to cell-autonomously regulate SSNs polarization and the coordinated dendritic outgrowth toward TCA^72,73^. Because we recently showed that CDKL5 loss reduces mGluR5 expression and function in the S1 cortex^29^, our current results prompt for further studies assessing the role of this glutamatergic receptor class in the alterations of the barrel system produced by the lack of CDKL5.

Barrel map formation is the result of a coordinated crosstalk between presynaptic TC inputs and L4 SSNs^74,75^, a circuit that we show to be severely affected in juvenile Cdkl5 KO animals. Our data show that the reduced number of TCA presynaptic boutons unlikely depends on difference in cellular density in the VPM nucleus nor to major alterations in of TCA, beside a slight decrease of their retrograde transport efficiency. In contrast, we found that SSNs dendrites in L4, which must receive highly organized TC inputs to promote tactile sensory processing in the BC^38^, show profound defects of their arborization and polarization in juvenile Cdkl5 KO mice, defects that can be caused by the recognized defects of microtubule-dependent anterograde trafficking in dendrites lacking CDKL5^76^. Moreover, in line both with our previous observation in S1 of adult Cdkl5 KO mice^47^ and with the role of CDKL5 in the postsynaptic compartment, we found that the maturation of spines decorating SSNs dendrites is defective in juvenile Cdkl5 KO mice. It is thus possible to conceive that such altered dendritic spines structure contributes to the reduction of TC synaptic contact that we report in L4 of the BC in Cdkl5 KO mice.

We disclose for the first time that mapping of tactile stimuli in the BC is aberrant in juvenile Cdkl5 KO, a defect that parallels atypical tactile responses shown by these animals during the first 15 postnatal days. Despite a highly disorganized and dysfunctional BC, the magnitude of evoked responses in awake juvenile Cdkl5 KO mice was exaggerated and not segregated within the limits of the corresponding C2 barrel when single-whisker deflection was applied. Interestingly, the orientation of SSNs dendritic processes toward the barrel hollow is necessary for bifacial cortical mapping of whiskers representation. In contrast, abnormal dendritic polarization in SSNs, as we observed in Cdkl5 mutants, results in impaired whiskers stimuli segregation that leads to an overlap with the responses from neighboring-whiskers^46,77^. Thus, our data provides first evidence that an ASD-related gene is important early during development for the morphological requirements of SNNs to promote correct cortical mapping of sensory stimuli. Moreover, although further studies are needed to clarify how the connectivity defects we found in juvenile Cdkl5 KO mice result in hypoactivation of the BC at resting conditions and to hyperactivation following whisker stimulation, it is feasible to hypothesize that early-onset dysfunctional tuning of sensory inputs leads to maladaptations in adult Cdkl5 mutant mice which will no longer show typical responses to sensory stimulation. We anticipate that our current findings, which appear in contrast with previous studies showing that cortical responses are reduced in adult Cdkl5 KO mice^27,29,49,78^, add instead novel information on the developmental trajectory of dysfunctional sensory tuning in Cdkl5 mutants.

Our data disclosed significant alterations of social behaviour in juvenile Cdkl5 KO mice starting from p21, a time when interaction with conspecifics begins^55^, and that that such early-onset atypical social responses were associated with autistic traits, such as repetitive or stereotyped behaviors (i.e., digging and jumping), as previously observed in adult Cdkl5 KO mice^27,79,80^. Because early cortical dysfunction leading to irregularities in tactile perception can, in turn adversely affect behavioral responses, we anticipated that the alterations in the BC of juvenile Cdkl5 KO mice promotes early defects in social interactions and autistic traits shown by these mutants. Intriguingly, our data show that neonatal replacement with a functional copy of CDKL5 restricted to the BC rescues both social behavior and autistic traits in Cdkl5 mutant mice, highlighting the existence of a causal link between aberrant development of whisker-to-barrel circuit and early-onset of autistic-like behavior in Cdkl5 KO mice. Although it will be of pivotal importance to assess in the future thalamic contribution, our data provide evidence that early CDKL5 activity within the BC circuitry is crucial for somatosensory perception, sociality and emotions, and ASD-like behaviors, a cortical area that should be taken into future consideration for targeting the autistic-traits associated with CDD.

In sum, by understanding the mechanisms through which ASD-related genes operate in crucial developmental periods, as we have attempted here with CDKL5, can foster the comprehension of autism-related aberrant sensory processing and improve the temporal tuning of personalized intervention plans in affected individuals.

## Supplementary materials

### MATERIALS AND METHODS

#### Animals

Animal care and experimental procedures were performed in accordance with European Community Council Directive 86/609/EEC for care and use of experimental animals and with protocols approved by the Italian Minister for Scientific Research (authorization D.M. n◦38/2020-PR 16/1/2020) and the Bioethics Committee of the University of Torino. Generation of the Cdkl5 line has been described previously^1^. Experiments were carried out on juvenile (p15-p25) Cdkl5 knockout (KO) male mice. Wild-type (WT) littermates were used for all experiments. After weaning, mice were housed together with their littermates in groups of 3-4 animals per cage and kept on a regular 12 h light/dark cycle (7:00-19:00 light period) and temperature-controlled environment (21 ± 2^◦^C). Food and water were available *ad libitum*. To maximally reduce the number of animals used in this study, mice were subjected to multiple behavioural tests, always starting with less invasive ones. All experiments were performed by an operator who was blinded to animal genotype or treatment.

### Histology and immunofluorescence

#### Coronal cortical sections

Juvenile mice (aged p15 for analyses shown in fig 1, 2, 3 and 4; aged p22 for fig 5; the number of animal used in each experiment is indicated in the result section) were deeply anesthetized by intraperitoneal injection (i.p.) of a mix of tiletamine/zolazepam (40 mg/kg) and xilazine (4-5 mg/kg) and rapidly perfused with phosphate buffered saline (PBS) solution 0.01M for 2 min, followed by a fixative solution consisting of 4% paraformaldehyde (PFA) in 0.01M phosphate buffer (PB) for 30 min. The brains were then removed, post-fixed overnight in 4% PFA at 4°C, and subsequently cryoprotected by immersing in 15% and 30% sucrose solution at 4°C. Free-floating 30 µm coronal sections were cut using a cryostat (Leica Biosystems), and stored at −20°C in a cryoprotectant solution containing 30% ethylene glycol and 25% glycerol until use (see^2^).

#### Flattened cortex sections

p15 mice were perfused as previously mentioned. Perfused brains were manually splitted in two hemispheres with a razor blade and, by using a small spatula, subcortical structures such as striatum, thalamus and hippocampus were removed to isolate the cortical tissue^3^. Subsequently, the resulting hemispheres were placed between two glass slides, filled with 4% PFA, and a weight was placed on top of it in order to make the hemispheres 10-20% thinner than the regular cortices. After 12h, at 4°C, the flattened cortices were washed (3x10min) with 0.1M PBS, cryoprotected and cut in 50 µm tangential sections.

#### Immunohistochemistry and cell analyses

Immunolabelling was conducted as in^4^. Briefly, sections were washed three times with 0.01M PBS (10 min) at room temperature and then permeabilized in 0.01M PBS solution containing 0.5% Triton X-100 in PBS and 5% normal donkey serum (NDS) for 60 min at room temperature. Subsequently, sections were incubated overnight at 4°C with the following primary antibodies: rabbit anti- Homer1bc (1:500, Synaptic System, 160 023); guinea pig anti-VGluT2 (1:1000, Millipore, 2251); guinea pig anti-VGluT1 (1:1000, Millipore, 5905); rabbit anti-c-fos (1:500, Cell Signaling Technology, 2250S); rabbit anti-GFP (1:1000, Invitrogen, A11122). The following day, sections were washed (3x10 min) with 0.01M PBS and incubated with suitable fluorescent secondary antibodies (1:1000; Jackson ImmunoResearch, West Grove, PA, USA) for 1 h at room temperature. Sections were then washed (3x10 min) with 0.01 M PBS, mounted onto glass slides with Dako fluorescence mounting medium (Dako Italia, Italy) and counterstained with the fluorescent nuclear dye DAPI (Sigma-Aldrich). Immunofluorescence images were acquired using a LSM 900 confocal microscope (Zeiss) and images were processed with ImageJ (NIH) or Photoshop CS6 (Adobe Systems). For puncta analysis, stacks of five optical sections (0.5 µm Z- step size) were acquired from L4 BC using a 63X objective (1 Airy unit). Synaptic puncta quantification and juxtaposition was performed by a custom written macro in ImageJ. For c-Fos^+^ cell analysis, stacks of eight optical sections (1 µm Z- step size) of BC were acquired using a 20X objective. Digital boxes spanning from the pial surface to the corpus callosum were superimposed at matched locations on each coronal section of the BC and divided into 10 equally sized sampling areas (bins; LI: bin 1; L2/3: bins 2–3; L4: bins 4–5; L5: bins 6–7; L6: bins 8–10). c-Fos^+^ immunopositive cells in each bin were manually counted as in^5^, while arbitrary intensity values from c-Fos^+^ cells were obtained using a dedicated ImageJ tool (integrative density) to analyse Z-stack projected images (Sum value). For c-Fos^+^ cell analysis following the novelty exposure test, flattened cortices were acquired using a 40X objective. Neurons positive for c-Fos were manually counted within the principal C2 barrel. C-Fos intensity analysis across C2 and adjacent barrels was performed as in^6^. In brief, C2-barrel was placed at the center of the image and cropped to include B2 and D2 neighbouring barrels. Following background subtraction, a plot profile of pixel intensities across the B2-C2-D2 barrels was generated with the Image J software (NIH, USA). To compare intensity profiles irrespective of genotype-associated differences in c-Fos signal intensity, intensities were binned and normalized to the mean intensity of C2.

### Adeno-associated virus (AAV)injections in the barrel cortex

Intracerebral injection procedures were performed as described by^7^, with minor changes. Briefly, hypothermic anesthesia of newborn (p0) pups was induced by placement on a cold aluminum foil and kept on ice for 5 min. Proof of anesthesia before injections were the observation of the mouse skin colour changing from pink to purple, and the absence of movement after gentle squeezing of the paws. Next, neonatal mice were placed in a stereotaxic neonatal mouse adaptor (Stoelting Co.) and the AAVs delivered using a glass micropipette attached to a nanoliter injector (Nanoject, Dummond Scientific Company) at a rate 23 nL/min. To target L4 of the BC, the following coordinates (in mm, measured from lambda and the pial surface) were used: AP: 1.5; ML: ±1.8; DV: 0.2. For TCA tracing, 50 nl at 1*10^12^ viral particles of retrograde AAV-hSyn-EGFP (Addgene 50465-AAVrg) were unilaterally infused. For sparse infection of neurons in the BC, 25 nl of AAV-hSyn-GFP (Addgene 50465-AAV1) at 1*10^11^ viral particles were bilaterally injected. For CDKL5 re-expression, 100 nL of AAVPHP.B_CDKL5 at 1*10^12^ viral particles were bilaterally injected as in^8^. Following injection, pups were placed on a warming pad for at least 10 min before being returned to the home cage.

### Neuronal morphology

To analyze retrogradely labeled TCA bundles, coronal sections were processed for immunofluorescence EGFP-signal amplification using an anti-GFP antibody. Images of the internal capsule and the dorsal striatum were captured from coronal sections using a 20X objective lens. A 100 µm x 100 µm region of interest (ROI) was selected in the dorsal striatum, and EGFP^+^ fibers were manually counted within this area. To normalize the EGFP expression for viral infection efficiency in injected animals, a ROI spanning the infection site was placed in the BC, and the integrated density of EGFP labeling was calculated using ImageJ software.

The dendritic polarization analysis of AAV-hSyn-GFP expressing L4 spiny stellate neurons (SSNs) was performed as in^9^. Following GFP signal amplification by immunolabelling with an anti-GFP antibody, images of single SSNs located at the barrel edge were acquired with the confocal microscope in flattened cortices using a 40X objective. Each dendrite was then traced with Image J and their length and arborization pattern was quantified using the orientation bias index (OBI). This index is defined as the ratio of inner dendrite length to total dendrite length. The average OBI of eight SSNs from the same animal was used as the OBI of the animal. Sholl analysis was performed as in^10^. Following dendrites tracing with the NeuronJ Plugin of ImageJ, concentric circles of 10 μm intervals were placed over GFP^+^ SSNs with the circles center positioned in the middle of the soma. Intersections of different dendritic orders and circles were then counted.

Dendritic spines decorating second order dendritic segments belonging to GFP^+^ SSN were acquired in tangential sections of flattened cortices with the use of the 40X objective lens of the confocal microscope. Spine number and size were analyzed as in^11,12^. Briefly, with ImageJ software both neck-length and head-diameter size were measured manually and employed to calculate neck/head ratios to categorize spine types as follows: mature spines, including mushroom (neck/head ratio < 1) and immature spines (neck/head ratio > 1). Mean spine density was measured as the number of spines per dendritic length unit (µm) with an independent replicate (animal) used as the sampling unit.

### Base scope ISH procedure

CDKL5 mRNA detection was performed as in^13^. Briefly, following PFA perfusion, brains from p25 mice were collected submerged in 4% PFA in 0.01M PBS for 24 h at 4 °C. Brains were then kept in sucrose solution (30%) at 4°C until sunk. Brains were then cut with a cryostat into 15 µm-thick coronal sections which were serially collected on glasses. The in-situ hybridization (ISH) for the CDKL5 RNA was performed with the Base Scope® technology (Biotechne) following the manufacturer’s protocol using a 1ZZ probe designed on the CDKL5 exon 4.

### Immunoblotting

#### Cortical extract preparation

Mice were sacrificed at p15 by decapitation, and the primary somatosensory cortices were dissected and homogenized in ice-cold RIPA buffer (NP-40 1%, Na deoxycholate 0,25%, EDTA 0.5M, NaCl 5M, Tris pH8 1M, SDS 10%, 1mM Na-Orthovanadate, 1 mM DTT, 0.5 mM PMSF, 50 mM NaF and protease inhibitors (SIGMAFAST™ Protease Inhibitor Cocktail Tablets, EDTA-Free). The lysates were then centrifuged at 13.000 g for 20 min at 4°C, the supernatant collected, and protein concentration measured using the BioRad protein assay method (BioRad).

#### Synaptosomal fraction preparation

Cortices from juvenile mice were homogenized in ice-cold lysis buffer (0.32 M sucrose, HEPES 1X at pH 7.4, 1 mM EGTA, 1mM Na-Orthovanadate, 1 mM DTT, phenylmethylsulphonyl fluoride and 1 mM sodium fluoride and protease inhibitors) (SIGMAFAST™ Protease Inhibitor Cocktail Tablets, EDTA-Free), using a glass Teflon tissue grinder. The homogenates were centrifuged at 1000 g for 10 min at 4°C. After discarding the nuclear pellet, the supernatant was centrifuged at 12,500 g for 20 min at 4°C. The P1 fraction was then washed with the same initial volume of lysis buffer and underwent further spin (20 min; 12,500 g). The pellet obtained, the crude cortical synaptosomal fraction (P2), was resuspended in 400 μl of RIPA buffer and stored at -80°C. Protein concentration of the synaptosomal fraction was determined with the Bio-Rad protein assay kit.

#### Western blotting

Proteins (30-50 mg) were mixed with Laemmli buffer 2x (375 mM Tris pH = 6.8, 12%SDS, 60% glycerol, 600 mM DTT, 0.06% bromphenol blue), heated to 90° C for 5min and then separated on SDS-PAGE gels as in^29^. Proteins were transferred to a PVDF membrane. The membranes were blocked for 1 h with BSA 5% or 1% milk in 1x TBS with 1% Tween (TBST) and incubated with the primary antibody overnight at 4°C. After washes (3x10 min) with TBST, the membranes were incubated with the appropriate secondary antibody for 1 h at room temperature. For total-protein assessment, the membranes incubated with the phospho-specific primary antibody were stripped with stripping buffer containing 2-mercaptoethanol, 1% SDS, and 62.5 mM Tris-HCl, pH 6.8, at 37°C for 30 min and reprobed with the relative total-protein antibody. The protein amount was normalized relative to the optical density of vinculin or β-actin. The following primary rabbit antibodies were used: NR1 (1:500, Invitrogen, PA5-85751); NR2A (1:1000, Invitrogen, PA5-35377); NR2B (1:1000,

Invitrogen, 71-8600); CamKII (1:1000, Millipore, 05-532); pCamKII (1:1000, Abcam, ab34703); AKT (1:1000, Cell Signaling, 4691P); pAKT (S473) (1:1000, Cell Signaling, 9271); ERK1/2 (1:1000, Cell Signaling, 9194); pERK (T202, T204) (1:1000, Cell Signaling, 4370); β-actin (1:1000, Cell Signaling, 4970); Vinculin (1:1000, Abcam, ab219649). The secondary antibody was a donkey anti-rabbit HRP (1:5000, Thermo Scientific, A16035). The chemiluminescent signal was visualized using Clarity™ Western ECL Blotting Substrates (Bio-Rad) and acquired with Bio-Rad ChemiDocTM Imager (Bio-Rad) and quantified with Image J software (NIH). Protein levels are displayed as fold change.

### Electrophysiology

#### Slices preparation

Juvenile (p15) mice were anesthetized by inhalation of isoflurane (Merial Italia) and sacrificed by decapitation. The brain was quickly extracted under hypothermic conditions and submerged in ice-cold cutting solution. For slices preparation, the cutting solution was composed of (in mmol/l): 70 sucrose, 80 NaCl, 2.5 KCl, 26 NaHCO3,15 glucose, 7 MgCl2·6 H2O, 1 CaCl2·2H2O, 1.25 NaH2PO4·H2O at pH 7.4 by saturation with 95% O2, 5% CO2. The region of interest was manually dissected by using a razor blade and blocked on the stage of a Microslicer DTK-1000 vibratome (Dosaka), using cyanoacrylate glue. 350 µm thick coronal slices of S1 cortex, including the barrel field, were cut, maintaining the tissue submerged in an ice-cold solution composed of (in mmol/l): 130 K gluconate, 15 KCl, 20 HEPES, 0.2 EGTA, 11 glucose. Slices were then transferred to a recovery chamber filled with the same solution continuously bubbled with 95% O2, 5% CO2, at room temperature (21–22 °C) for at least 1 h before starting the recording. The incubating solution used for the voltage-clamp recordings was composed of (in mmol/l): 125 NaCl, 2.5 KCl, 26 NaHCO3, 15 glucose, 1.3 MgCl2, 2.3 CaCl2·2 H2O e 1.25 NaH2PO4·H2O.

#### Voltage-clamp recordings

NMDAR activated currents were recorded in voltage clamp conditions and whole-cell configuration as in^14^. Briefly, pyramidal cells were visualized by means of an upright microscope (Axioskop 2 FS; Zeiss, Oberkochen, FRG) equipped with a ×60 water-immersion objective lens, differential-contrast optics, and a near-infrared charge-coupled device (CCD) camera. Slices were perfused with ACSF (continuously bubbled with 95% O2, 5% CO2) at a rate of about 1.5 ml/min. Patch pipettes were fabricated from thick-wall borosilicate glass capillaries (CEI GC 150-7.5; Harvard Apparatus, Edenbridge, UK) by means of a Sutter P-87 horizontal puller (Sutter Instruments, Novato, CA, USA). NMDAR-mediated currents were recorded by holding neurons at -70 mV and perfusing them with the NMDAR agonist, N-Methyl-D-aspartate, (NMDA, 50 µM). The external solution contained (in mM): 130 NaCl, 1.8 CaCl2, 10 HEPES, 10 glucose, 1.2 Glycine (pH 7.4). The internal solution contained (in mM): 90 CsCl, 20 TEACl, 10 glucose, 1 MgCl, 4 ATP, 0.5 GTP, 15 phosphocreatine (pH 7.4). These experiments were performed in the presence of the AMPA receptors blockers: 6,7- dinitroquinoxaline-2,3-dione, DNQX (20μM, Sigma-Aldrich). Tetrodotoxin (TTX 1 µM) was added to block voltage-gated Na^+^ channels. Current signals were acquired in gap-free modality with a PC interfaced to a Digidata 1322A interface (Axon Instruments) using the Clampex program of the pClamp 8.2 software package (Axon Instruments). Current signals were low-pass filtered at 5 kHz and digitized at 20 kHz.

### Intrinsic optical signal (IOS) imaging

#### Surgery

The surgery was performed, as previously described in^15^, under isoflurane anesthesia (3% for induction and 1% for maintenance) on juvenile male mice (p22-25). After being placed on a stereotaxic frame and head fixed using ear bars, local anesthesia was provided using subcutaneous lidocaine (2%) injection and eyes were protected with dexamethasone-based ointment (Tobradex, Alcon Novartis). The scalp was removed, and the skull carefully cleaned with saline. Next, a thin layer of cyanoacrylate was poured over the exposed skull to attach a custom-made metal ring (9 mm internal diameter) centered on the barrel cortex. When the glue dried off, a drop of transparent nail polish was spread over the area to ameliorate optical access. During the entire procedure body temperature was controlled using a heating pad and a rectal probe to maintain the animals’ body at 37°C. The animals were left for an hour to recover before the IOS recordings.

#### IOS recordings

IOS recordings were performed under isoflurane (1%) and chlorprothixene (1.25 mg/Kg, i.p.). Images were obtained using an Olympus microscope (BX50WI) under red light illumination provided by 8 red LEDs (625 nm, Knight Lites KSB1385-1P) attached to the objective (Zeiss Plan-NEOFLUAR 5x, NA 0.16) using a custom-made 3D printed LED holder controlled by an Arduino hardware (Arduino UNO). The animal was secured under the objective using a ring-shaped neodynium magnet (www.supermagnete.it, R-12-09-1.5-N) mounted on an Arduino-based 3D printed imaging chamber that also controls eye shutters and a thermostated heating pad. Visual stimulation was performed using Matlab Psychtoolbox and presented on a monitor placed 13 cm away from the eyes of the mouse: visual stimulus consisted of a sinusoidal wave grating presented in the binocular portion of the visual field with spatial frequency 0.03 cyc/deg, mean luminance 20 cd/m^2^ and contrast 90%. The stimulus consisted in the abrupt contrast reversal of a grating with a temporal frequency of 4 Hz for 1 s.

For whisker stimulation, we used piezoelectric bending actuators (PI P-615.30D) connected to a wire custom-made arm touching all whiskers. The stimulus was generated by a sweep generator (Wavetek model 164) connected to an amplifier (E-663 LVPZT-amplifier). Stimuli consisted of 1 mm back-and-forth deflections at 4Hz for 1 sec, on one side of the snout, 0.5 cm away from the face. Frames were acquired at 30 fps, with a resolution of 512 x 512 pixels. A total of 270 frames were captured for each trial: 30 before the stimulus as a baseline condition and 240 as post-stimulus. The signal was averaged for at least 30 trials and down sampled to 10 fps. Fluctuations of reflectance (R) for each pixel were computed as the normalized difference from the average baseline (ΔR/R). For each recording, an image representing the mean evoked response was computed by averaging frames between 0.5 to 2.5 sec after stimulation. The mean image was then low-pass filtered with a 2D average spatial filter (30 pixels, 117 μm^2^ square kernel). To select the barrel cortex for further analysis, a region of interest (ROI) was automatically calculated on the mean image of the response by selecting the pixels in the lowest 20% ΔR/R of the range between the maximal and minimal intensity pixel^16^. To weaken background fluctuations a manually selected polygonal region of reference (ROR) was subtracted. The ROR was placed where no clear response, blood vessel artifact or irregularities of the skull were observed^17^. Mean evoked responses were quantitatively estimated as the average intensity inside the ROI.

### Innate sensorimotor reflexes

The examination of the development of motor and sensory performances was carried out from p3 to p15 as described^18^. The analysis of innate reflexes was combined with the investigation of developmental milestones including: eye opening, incision eruption, body weight, and ear detachment. Individual pups were behaviourally tested in the following order: surface righting test, cliff avoidance test, negative geotaxis test, grasp test, whisker stimulation test, and tactile stimulation test.

<u>Surface righting reflex:</u> mice were placed in the supine position, held for 5 sec and then released. The latencies required to roll over and have all four paws in contact with the surface were recorded. The maximum time allowed per trial for righting was 60 sec. The score was considered absent (0) if the pup lied on its back for 60 sec, 1 if it turned in more than 1 sec, and 2 if it turned in less than 1 sec. Mice were tested in 2 trials for each daily session and the mean score was calculated.

<u>Cliff avoidance test:</u> animals were placed on an edge of elevated platform with both forepaws and nose over the cliff. The mouse’s ability to crawl back and turn away from the cliff was scored: (0) pup fell down, (1) pup turned less than 90°, (2) turned 180°. Each pup was tested 2 times in each experimental session and the mean score was calculated.

<u>Negative geotaxis:</u> pups were placed on an inclined plywood surface (45° angled, 20×20 cm) and observed for up to 30 seconds. The score was considered (0) when pups did not turn around and fell down, (1) < 90°, (2) 90°< x<180° turn and (3) 180° turn. Each pup was given 2 trials each test day and the mean score was calculated.

<u>Grasp test:</u> pups were placed in a glass Petri dish and the experimenter gently applied a little pressure against the palmar surface of a forepaw with a thin rod and the palmar grasp reflex was recorded. This reflex is characterized by closing of the digits in response to an object stroking the palmar surface. The ability to perform this reflex was assigned as: absent (0 points), crooked digits but not grasped (0.5 point), or present (1 point) in each of the forelimbs separately. The score of the grasping reaction was: absent (0 points), grasp but not sustained (0.5 point), or sustained (1 point) in each of the forelimbs separately. Each pup was examined in 2 consecutive trials and the mean of this score was calculated.

<u>Whisker stimulation test:</u> this test was performed to assess behavioral reaction, such as twitching responses and head moving reactions, after unilateral whiskers stimulation with a thin stainless-steel rod (0.5 mm in diameter) for 20 sec without touching the inter-vibrissal fur or the snout skin. A score of 1 was assigned when a reaction was shown, and 0 when it was absent.

<u>Tactile stimulation test:</u> pups were kept in a glass Petri dish for 1 min, following the whisker stimulation test, and several points along the body, including the top of the head, medial back, left and right forepaws, hind paws, and the base of the tail (total 7 points) were manually poked 5 times with a thin stainless-steel rod (0.5 mm diameter). The reaction of mice, for each stimulated part of the body, was measured. A score of 1 was assigned when a reaction was shown, and 0 when it was absent.

### ASD-related behavioral tests

The behavioral studies were carried out in the morning on mice aged p21-25. Mice were habituated to the test room for at least 1 hour before each test and the behavior tests housing cleaned between each trial with 70% ethanol.

<u>Open field test:</u> exploratory behavior in a novel environment was assessed by a 20 min session within an open field arena (50cmL x 50cmW x40cmH) made of gray Plexiglas. Juvenile mice (p21-25) were placed in the center of the arena and then the videorecording using a digital camera was started. Locomotor activity (distance traveled and speed, resting time) in the entire arena were then analyzed using the Smart 3.0 software (Panlab).

<u>Juvenile play test:</u> juvenile social interactions were assessed in 21-22 day-old pairs of male mice from different litters as previously described^19^. Prior to the social interaction session, mice were individually housed for 1 hour in a standard laboratory cage, without access to food and water. After an hour of individualized housing, each mouse was placed alone in the play testing cage for a 10-min habituation period. The floor of the arena was covered with a thin and fresh layer (0.5 cm) of the same type of bedding that is used for the home cage. After mice were habituated, a pair of animals of the same sex was placed simultaneously in the testing arena and video recording immediately started. Two genotype combinations were assessed: WT/WT and KO/KO mice. Mice were recorded for the initial 10 min, when social interaction is active. Behavioral activities including anogenital sniffing, nose-to-nose sniffing, body sniffing, push crawl and follow, were manually evaluated by an investigator blinded to the genotypes.

<u>Three chamber sociability test:</u> the social interaction between mice was tested as described in^20^. Briefly, the testing apparatus consisted in a rectangular grey Plexiglas three chamber box (60 cm (L) x 40 cm (W) x 20 cm (H) equipped with dividing walls and doorways allowing access to each chamber. Age and sex matched animals were used for all tests. WT mice were used as stranger mice and were habituated to placement inside the wire cage. After 10 minutes of habituation in the central chamber, test mice were then allowed to explore the entire arena, with open access to both left and right chamber. After 10 min, during the social phase, the stranger was placed in one chamber while an empty round wire cup was placed in the other chamber. The test animal was allowed to freely explore the social apparatus for 10 min and show whether it prefers to interact with the inanimate object or with the stranger mouse. Time spent in each chamber, as well as locomotion, was calculated using automated software (Smart 3.0, Panlab). The sociability index (SI) was calculated as in^21^.

<u>Repetitive behavior:</u> mice were individually placed into a clean home-cage like environment lined with bedding. After 5 min of habituation, 10 min of activity was recorded. The number of repetitive behavior, defined as grooming, digging and jumping was scored manually using a stopwatch as described^22^.

### Novelty exposure test

This test was performed in p21 Cdkl5 and WT littermates. The day before the experiment, mice were habituated to the testing room and arena. Afterwards, all the whiskers of the snout were completely trimmed except the C2 whisker on the left whisker pad. The day after, the mouse was placed in the testing arena filled with objects with different textures. Mice were free to investigate novel objects for 1 hour while their track was recorded. Immediately after, mice were perfused, the hemisphere flattened, and the tissue processed for c-Fos staining as described in^23^.

### Statistics

Statistical analysis was performed using Prism software (Graphpad). Statistical values including the n, statistical test and significance are reported in the figure legends. The data were analyzed by: (i) one- or two-way analysis of variance (ANOVA) followed by the Bonferroni test, if differences were found; (ii) unpaired t-test; (iii) Mann–Whitney test. All data are reported as mean ± SEM and considered to be statistically significant when p < 0.05. In each graph circles/dots represent individual values. From all analyses, outlier values were excluded according to the ‘‘identify outliers” function (Method: ROUT, Q= 2%) present in Prism software. Power analysis of the statistical tests was performed using G Power 3.1.9.2 (RRID:SCR_013726).

## SUPPLEMENTARY FIGURES

**Supplementary Figure 1:**
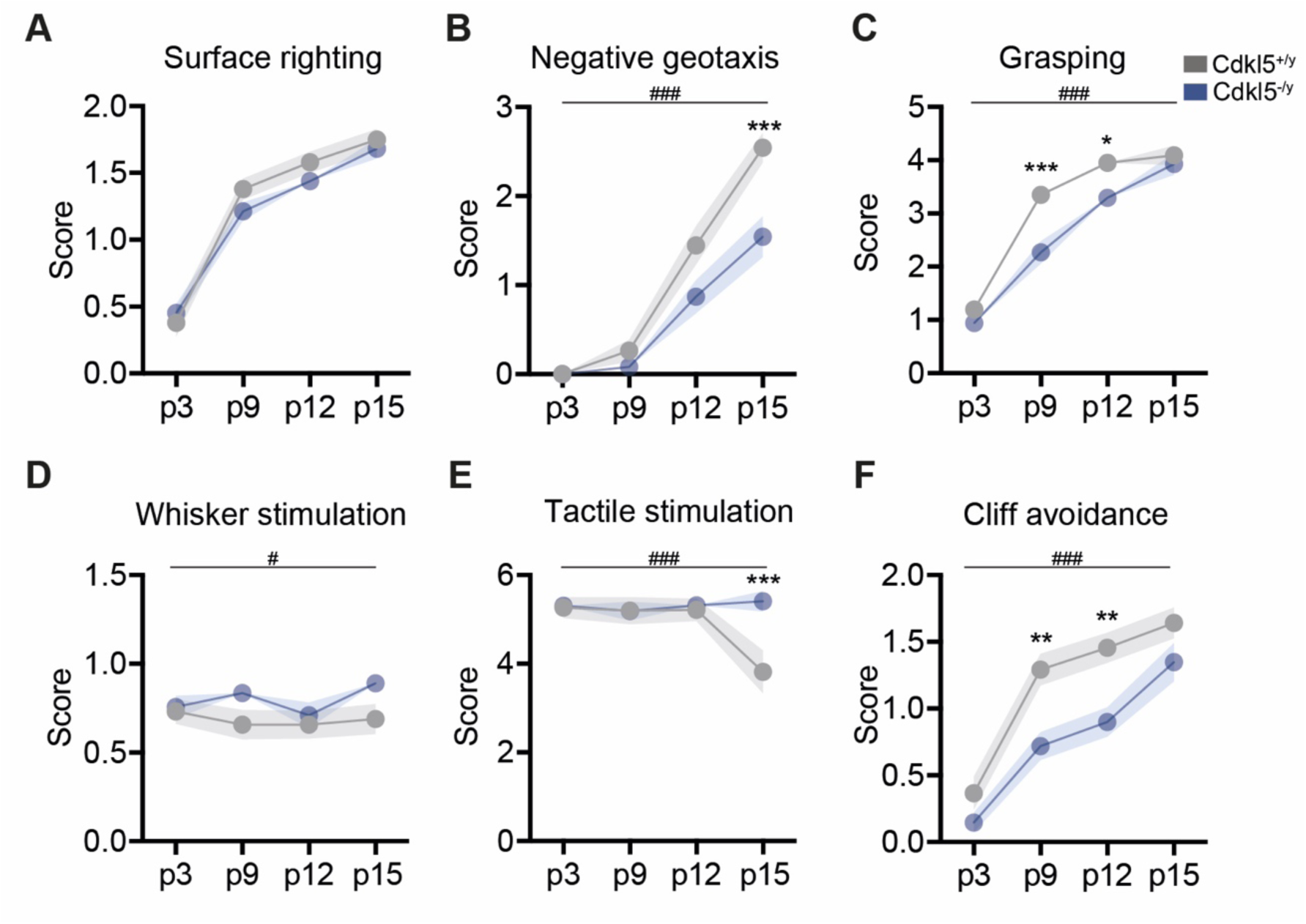
Cdkl5 KO mice show early atypical innate sensory-motor reflexes. (A-C) Motor performance in Cdkl5^+/y^ and Cdkl5^-/y^ pups. The parameters scored are: the latency for righting their body for the surface righting test (A), the latency for body rotation for the negative geotaxis test (B) and the combination of grasp reflex and grasping strength for the grasp test (C). (D-F) Sensory performance in Cdkl5^+/y^ and Cdkl5^-/y^ pups. The parameters scored are: the motor response as a reaction to 30 times whisker stimulation for the whisker stimulation test (D), the twitching response to body stimulation for the tactile stimulation test (E) and the latency to avoid the cliff for the cliff avoidance test (F). Data are expressed as mean ± SEM. Cdkl5^+/y^ = 25; Cdkl5^-/y^ = 30. Statistical analysis: 2-way ANOVA, genotype effect #p < 0.05, ### p < 0.001. Bonferroni’s post-hoc test, *p < 0.05, **p < 0.01, *** p < 0.001.

**Supplementary Figure 2.**
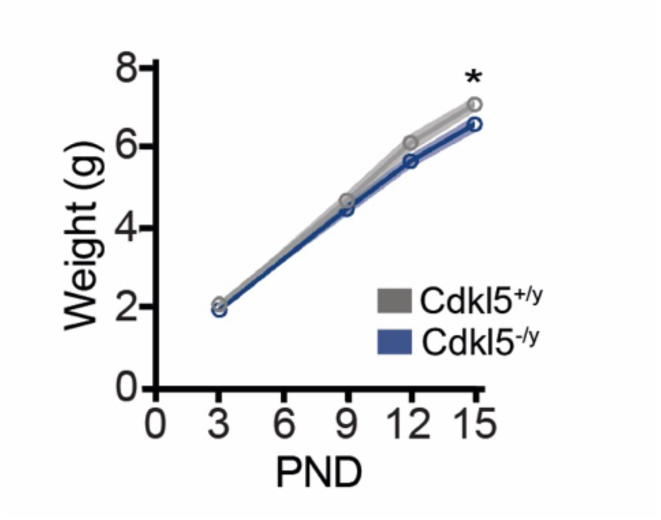
Cdkl5^-/y^ mice show normal body growth until P15 compared to Cdkl5^+/y^ littermates. n = 25-29. Statistical analysis: 2-way ANOVA, Bonferroni’s post-hoc test, *p < 0.05.

**Supplementary Figure 3.**
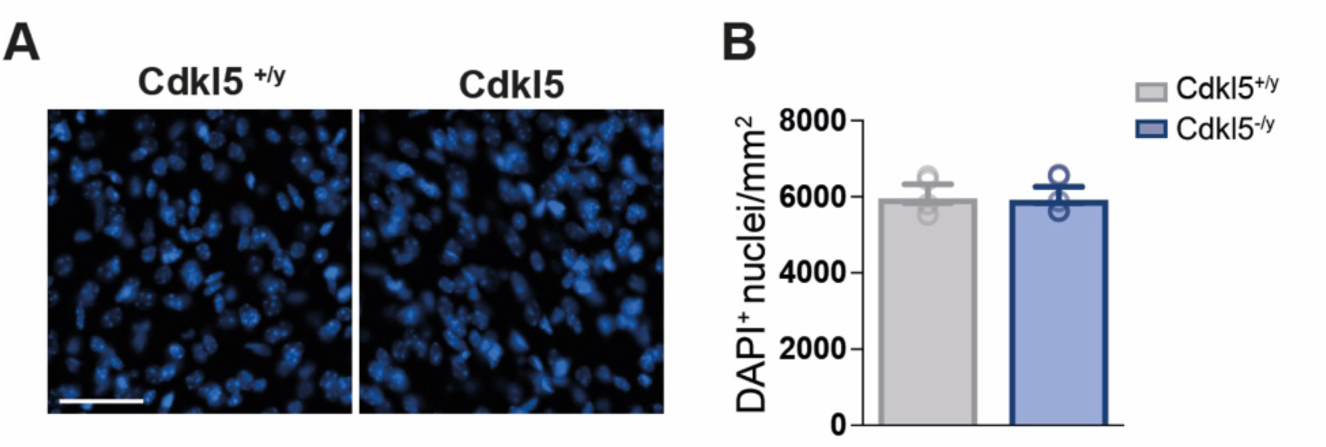
A normal VPM nuclear density in juvenile Cdkl5^-/y^ mice. (A) Micrographs showing the DAPI staining in the VPM of Cdkl5^+/y^ and Cdkl5^-/y^ mice and histogram (B) showing the quantification of nuclei number (n = 4). Statistical analysis: t-test.

